# MGMG: Cell Morphology-Guided Molecule Generation for Drug Discovery

**DOI:** 10.1101/2025.07.11.664424

**Authors:** Qiaosi Tang, Daoyun Ding, Xiaoyong Yuan, Gustavo Seabra, Peter A Ramdhan, Chi-Yuan Liu, My T. Thai, Chenglong Li, Hendrik Luesch, Yanjun Li

**Author notes:** Correspondence: Yanjun Li.

## Abstract

Designing novel molecules with desired bioactivity remains a fundamental challenge in drug discovery. Most molecular design methods follow target-based drug discovery paradigms that rely on well-defined drug targets, thereby limiting their applicability to diseases lacking known targets or reference compounds. Here we introduce Morphology-Guided Molecule Generation (MGMG), a phenotypic drug discovery–oriented approach that integrates cellular morphological profiles from compound treatments with molecular textual descriptions without requiring molecular target information. Cell morphology offers the guidance on desired bioactivity-relevant phenotypic effects, while textual descriptions provide direct and interpretable cues about molecular structure. Leveraging complementary structural and bioactivity context, MGMG significantly enhances molecule generation performance, especially in scenarios where textual descriptions are under-informative or morphological signals are weak. MGMG can also be applied to genetic perturbations, enabling activator design from gene overexpression morphology without requiring knowledge of reference compound structure. In addition, in silico docking demonstrates that MGMG-generated molecules, despite lacking target information, exhibit binding affinities comparable to reference compounds, preserving key interactions while introducing structural diversity. Overall, MGMG jointly utilizes morphological and textual description inputs to guide molecule generation, enabling diverse, bioactivity-aware compound design in a target-agnostic fashion.

## Introduction

Drug discovery is a challenging and time-consuming process; modern artificial intelligence-aided drug design (AIDD) methods aim to accelerate it by enabling the automated identification of drug candidates^1^. Most AI-based molecular design methods adhere to the principles of target-based drug discovery (TDD), relying on well-defined biological molecular targets and binding pockets for structure-based drug design^2–6^, or on known active molecule binders as reference compounds for ligand-based molecular generation^7–9^. Consequently, these methods have limited applicability for diseases lacking validated targets or existing reference compounds.

In contrast to TDD, phenotypic drug discovery (PDD) does not depend on knowledge of specific drug targets or hypotheses about their roles in diseases. Instead, it relies on the chemical interrogation of disease-relevant biological systems in a molecular target-agnostic fashion and has successfully identified many of the first-in-class drugs approved by the US Food and Drug Administration (FDA) over the past decades^10, 11^. High-content imaging (HCI), enabled by automated microscopy and image analysis, has paved the way for large-scale, high-throughput drug screening in PDD^12^. For example, the Cell Painting assay^13, 14^, a prominent and low-cost HCI technique, provides a comprehensive view of the cellular state by staining major cellular components and producing morphological profiles as phenotypic descriptors. These morphological profiles have led to a comprehensive understanding of cellular behavior under chemical perturbations^13^. The target-agnostic nature and the ability to provide significant insights into biological responses make morphological profiling well-suitable for addressing the challenges faced by TDD-oriented *de novo* molecule design methods.

While promising, only two previous studies have explored morphological profiling in the task of molecule design^15, 16^, and both exhibit several limitations. First, because many different molecules can induce similar morphological effects, relying solely on cell morphology data for molecule generation often lacks specificity and control over the desired molecular structure or scaffold. Particularly in PDD, where drug target information is not available as a generation condition, the absence of additional guidance beyond the morphological data can lead to the omission of essential molecular properties, such as synthesizability and drug-likeness^15, 16^, limiting the applicability of the generated molecules. Furthermore, generating molecules based on cell images from weak or inactive perturbations, which induce only subtle or minimal morphological changes, poses challenges in extracting clear, actionable signals for molecule generation^15^.

In this work, we introduce **MGMG** (**M**orphology-**G**uided **M**olecule **G**eneration), an AI-driven molecule design approach tailored for PDD, which addresses these challenges by enabling morphology-informed bioactivity awareness and structural and property control within a unified framework. To achieve this, we propose integrating cellular morphological data from compound treatments with molecular textual descriptions to jointly guide molecule design, without requiring molecular target information. Complementary to morphology, molecular textual descriptions offer a high degree of interpretability and have proven effective in conveying open-ended chemical information about molecular structures and properties across diverse drug discovery tasks^17–20^. Meanwhile, in medicinal chemistry, the availability of domain knowledge and heuristic “rules of thumb” are essential factors guiding real-world molecule discovery campaigns. Thus, driven by its unique advantages, we leverage molecular textual descriptions to embed domain knowledge of the desired structural and physicochemical attributes, complementing the guidance provided by morphological data. Through pretraining and contrastive learning paradigms, MGMG bridges the two modalities and their guidance signals by aligning them into a unified latent space and capturing their interactions. This combination enables molecule design with simultaneous bioactivity awareness, specificity, and structural control, overcoming the limitations of methods that rely solely on cell morphology. In parallel, unlike molecule captioning and text-based molecule generation approaches, our method provides explicit guidance on molecular biological effects and mechanisms of action, encoded through cellular morphological profiles. These aspects are challenging to capture precisely and thoroughly in textual descriptions, whereas cell morphology offers comprehensive, low-level phenotypic representations that are crucial for drug discovery and design.

Through experiments, we demonstrate that the proposed approach effectively enhances specificity and control in molecule generation, achieving a higher recovery rate while maintaining a degree of structural diversity. We further show that the two input modalities compensate for each other, with morphological profiles improving molecule generation when textual descriptions are under-informative, and textual descriptions enhancing performance when morphological signals are weak. Our model can be extended from chemical to genetic perturbations, enabling activator design based on morphological profiles from gene overexpression, without requiring structural guidance from reference activators, even in cases of weak perturbations. We also explore whether the molecules generated in this target-agnostic manner can potentially bind to known protein targets. Our results show that the generated molecules can retain key interactions with target protein residues despite the input lacking explicit structural knowledge of the target. In some cases, structural variations introduced by MGMG enable the molecules to explore the binding pocket and access novel chemical space. In summary, we present MGMG as a versatile framework for the design of diverse and high-quality molecules with bioactivity context in a disease target-agnostic fashion.

## Results

### Overview of MGMG

We introduce MGMG, a PDD-oriented small molecule generation framework that leverages both desired cellular morphological profiles and molecular textual descriptions to jointly guide the design of novel compounds (Fig. 1a). Cell morphology serves as a functional readout of chemical perturbations, capturing the biological impact of a designed compound at the phenotypic level. In contrast, textual descriptions convey structural and physicochemical information, reflecting the compound’s properties at the molecular level. MGMG integrates these complementary modalities to enable molecule generation that is both biologically informed and structurally guided.

**Fig. 1:**
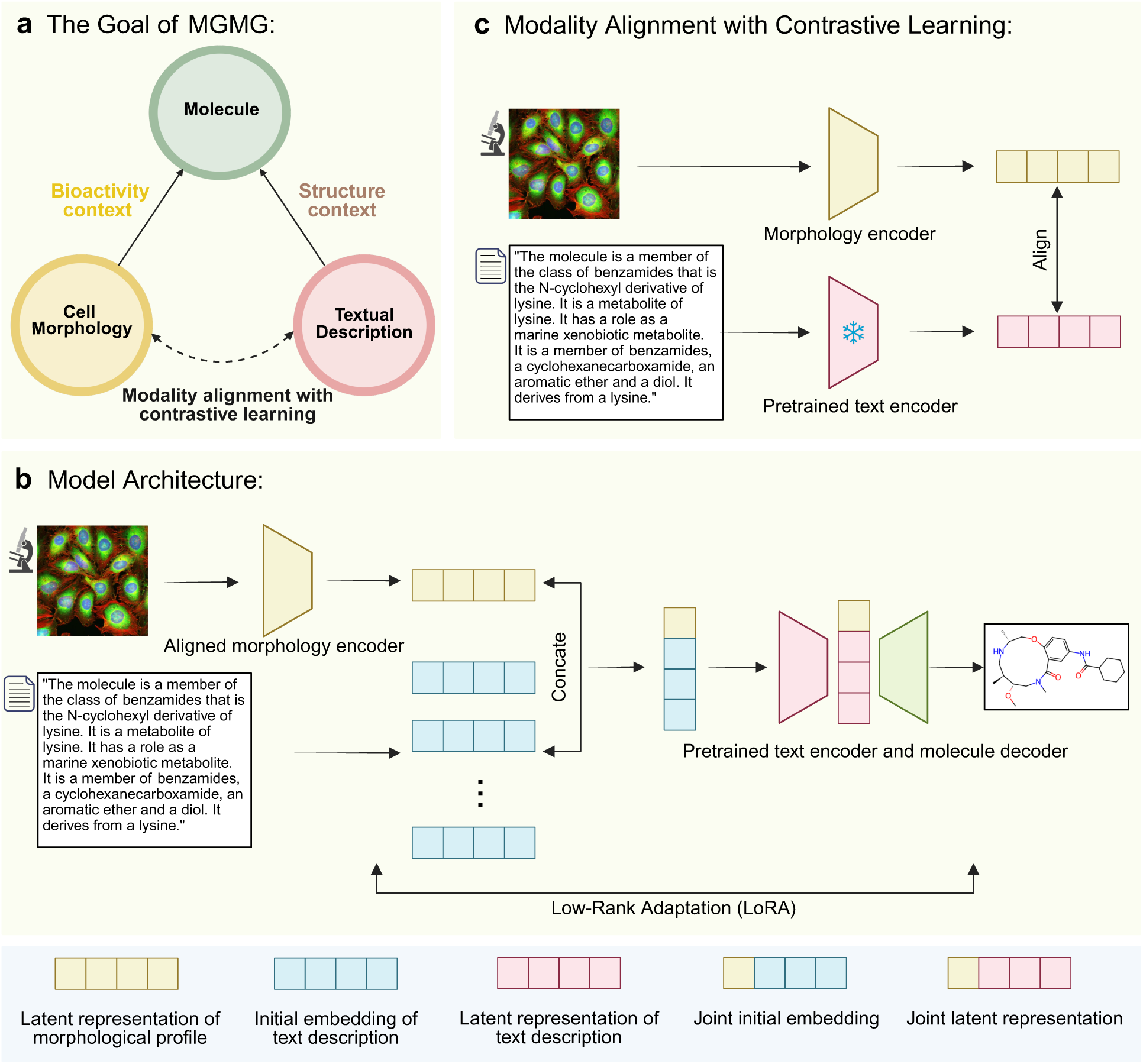
Schematic of MGMG. **a** The overarching goal of MGMG is to design novel small molecules guided by cell morphology–informed bioactivity signals and molecular textual descriptions of a reference compound, without requiring prior knowledge of the molecular target. **b** Within an encoder-decoder Transformer framework, the complementary modalities are integrated using an early fusion approach and jointly guide molecule generation in the SELFIES format. **c** The contrastive learning strategy is utilized to align cell morphology profiles and textual descriptions to establish a unified latent representation space.

MGMG is built on an encoder-decoder Transformer architecture (Fig. 1b). The encoder comprises two branches: a cell morphology branch that processes the cellular morphological profile induced by the reference compound, and a textual description branch that extracts semantic representations from the reference compound’s textual description. The disentangled design allows MGMG to integrate the powerful existing models pretrained on a single modality. Specifically, we utilize the BioT5 model^21^ pretrained on the large-scale text-molecule pairs as the base model for the text branch encoder to bridge the textual description associations and chemical structure. The morphology branch uses a multi-layer perceptron network to encode morphological profiles extracted by the bioimaging software CellProfiler^22^, which generates predefined biologically insightful features from raw Cell Painting images. These profiles have been successfully and widely used for characterizing compound mechanisms of action in phenotypic drug discovery^23^. To establish a unified representation space between these two modalities, MGMG employs the contrastive learning strategy^24^ to align the morphology branch to the pre-trained text branch (Fig. 1c). After the alignment, the dual-modality inputs are integrated using an early fusion approach to generate a joint encoder representation, upon which a decoder autoregressively generates novel molecules in SELFIES format^25^.

To train MGMG, we first curated a dataset named Mol-Instructions-BBBC036v1, a structure–text–morphology dataset comprising 25,070 compounds, by pairing morphological profiles from BBBC036v1^26^ with molecular textual descriptions from PubChem^20, 27^ using the unique compound structures as the linking key. However, many of the original textual descriptions were under-informative with numerous distinct molecules sharing the same brief annotations, limiting their utility in molecule generation tasks. To enhance the quality and specificity of these descriptions, we employed the pre-trained molecular captioning model BioT5^21^ to generate synthetic, detailed textual descriptions for each compound. The resulting dataset, MolCaptioned-BBBC036v1, was first used for multimodal alignment between the morphology and text branch encoders, followed by end-to-end encoder-decoder training for the entire MGMG molecule generation framework. Given the relatively limited dataset size, the Low-Rank Adaptation of Large Language Models (LoRA) was adopted to mitigate overfitting^28^.

### MGMG generates molecules with desired chemical properties

To our best knowledge, MGMG is the first approach to integrate both morphological and textual descriptions for *de novo* molecule design. Accordingly, we compared MGMG with the state-of-the-art unimodal baselines to assess its performance. Among the two existing morphology-based approaches, we compared MGMG to CPMolGAN, a Generative Adversarial Networks (GAN)-based model that generates molecules using morphological profiles as input^15^. We did not include the GFlowNet-based model in our comparison, as it requires training a separate model for each target compound, which does not scale to the size of our training dataset or align with our evaluation setup^16^. In addition to morphology-based models, we also compared MGMG with text-based molecule generation models, such as MolT5^29^ and BioT5^21^. Notably, MGMG is a light model with only 14 million trainable parameters—just 5% of the model size of BioT5 or MolT5-base (Supplementary Fig. 1). Because molecules can be represented as biosequence-like structures (e.g., SMILES or SELFIES), we evaluated the generated outputs using natural language processing metrics such as BLEU^30^ and Levenshtein distance^31^ to measure the similarity between generated and reference SELFIES or SMILES. Results in Table 1 demonstrate that MGMG outperforms all baseline methods on both metrics. In addition, both MGMG and BioT5 achieve 100% validity in the generated molecules, underscoring the robustness and error-tolerance of using SELFIES representations for molecular generation.

**Table 1.** Properties of molecules generated by different models on MolCaptioned-BBBC036v1.

| Property | Details | MGMG | BioT5 | CPMolGAN | MolT5 |
| --- | --- | --- | --- | --- | --- |
| Reconstruction metrics | BLEU score↑ | **0.832±0.003** | 0.821±0.002 | 0.244±0.036 | 0.545±0.0005 |
|  | Levenshtein score↓ | **14.730±0.176** | 15.613±0.278 | 44.551±1.171 | 42.409±0.028 |
| Physiochemical properties | MW | 430.712±0.560 | 430.056±2.382 | 424.881±27.531 | 454.936±0.781 |
|  | MW R<sup>2</sup> to reference↑ | **0.848±0.005** | 0.831±0.008 | -0.056±0.277 | 0.282±0.004 |
|  | LogP | 2.956±0.035 | 2.946±0.026 | 3.817±1.929 | 3.643±0.004 |
|  | LogP R<sup>2</sup> to reference↑ | **0.713±0.007** | 0.689±0.010 | -0.002±0.082 | 0.120±0.004 |
|  | HBA | 5.466±0.042 | 5.469±0.058 | 5.083±2.234 | 4.895±0.010 |
|  | HBA R<sup>2</sup> to reference↑ | **0.765±0.012** | 0.744±0.003 | -0.053±0.108 | 0.287±0.009 |
|  | HBD | 1.777±0.012 | 1.784±0.037 | 1.673±0.628 | 1.720±0.008 |
|  | HBD R<sup>2</sup> to reference↑ | **0.752±0.013** | 0.740±0.006 | -0.056±0.086 | 0.136±0.008 |
| Drug-likeness | ChEMBL FCD↓ | **2.546±0.078** | 2.748±0.046 | 43.696±10.743 | 16.616±0.096 |
|  | QED↑ | **0.589±0.003** | 0.588±0.006 | 0.486±0.053 | 0.555±0.001 |
|  | %QED>0.5↑ | **0.714±0.009** | 0.710±0.006 | 0.462±0.135 | 0.639±0.004 |
|  | R<sup>2</sup> to reference↑ | **0.572±0.013** | 0.526±0.014 | -0.032±0.112 | 0.092±0.011 |
| Synthesizability | SA score↓ | **3.695±0.014** | 3.725±0.024 | 3.713±0.628 | 4.209±0.005 |
|  | %SA score<4.5↑ | **0.885±0.007** | 0.871±0.004 | 0.802±0.117 | 0.613±0.001 |
|  | R<sup>2</sup> to reference↑ | **0.536±0.008** | 0.508±0.014 | -0.060±0.054 | 0.129±0.004 |
| Diversity | %Unique Murcko Scaffold↑ | **0.748±0.003** | 0.743±0.005 | 0.595±0.260 | 0.456±0.004 |
| Validity | %Valid↑ | **1.00±0.00** | 1.00±0.00 | 0.841±0.087 | 0.928±0.003 |
| Uniqueness | %Unique↑ | **0.980±0.001** | 0.977±0.0002 | 0.732±0.291 | 0.663±0.002 |

Generating molecules with desired properties is crucial for drug discovery. Herein, we evaluated the molecules generated by MGMG and baseline methods based on a wide range of physicochemical, drug-likeness and synthesizability properties, even though these attributes were not monitored or optimized during the training process. Most molecules generated by MGMG adhere to the “Lipinski’s Rule of Five”^32^, with an average Molecular Weight (MW) <500 Da, Lipophilicity (LogP) <5, hydrogen bond acceptors (HBA) <10, and hydrogen bond donors (HBD) <5 (Table 1). We observed a high correlation between the physicochemical properties of MGMG-generated molecules and their corresponding reference compounds (Fig. 2a, Table 1), further demonstrating the model’s capability to accurately recover biologically relevant molecular structures. Supplementary Fig. 2 presents comparative examples of molecules generated by MGMG, BioT5, MolT5, and CPMolGAN. Overall, MGMG and BioT5 produced molecules structurally similar to the reference compound, with MGMG showing greater similarity. Molecules generated by MolT5 and CPMolGAN diverged more substantially from the reference compound. These observations were further supported by quantitative evaluations of drug-likeness and structural realism. MGMG generated molecules with the highest average drug-likeness (QED) scores among all models as well as the highest percentage of molecules exceeding a QED threshold of 0.5, indicating strong alignment with drug-likeness criteria. Moreover, the QED scores of MGMG-generated molecules showed the strongest correlation with those of the reference compounds, further highlighting its ability to preserve key drug-likeness attributes. In terms of structural realism, MGMG-generated molecules achieved the lowest Fréchet ChemNet Distance (FCD) scores^33^, indicating a closer resemblance to the distribution of known bioactive compounds. In addition to drug-likeness, we also examined the synthetic accessibility (SA) score, which estimates the ease of synthesis^34^. MGMG achieved the lowest average SA score and the highest percentage of molecules with an SA score below 4.5. It also exhibited the strongest correlation between the SA scores of generated molecules and their reference counterparts. These findings indicate that MGMG generates molecules with greater synthesizability than those produced by other models. Collectively, the strong performance across these criteria highlights MGMG’s ability to generate novel molecules with desired chemical properties. Detailed descriptions of these evaluation metrics are provided in the Methods section.

**Fig. 2:**
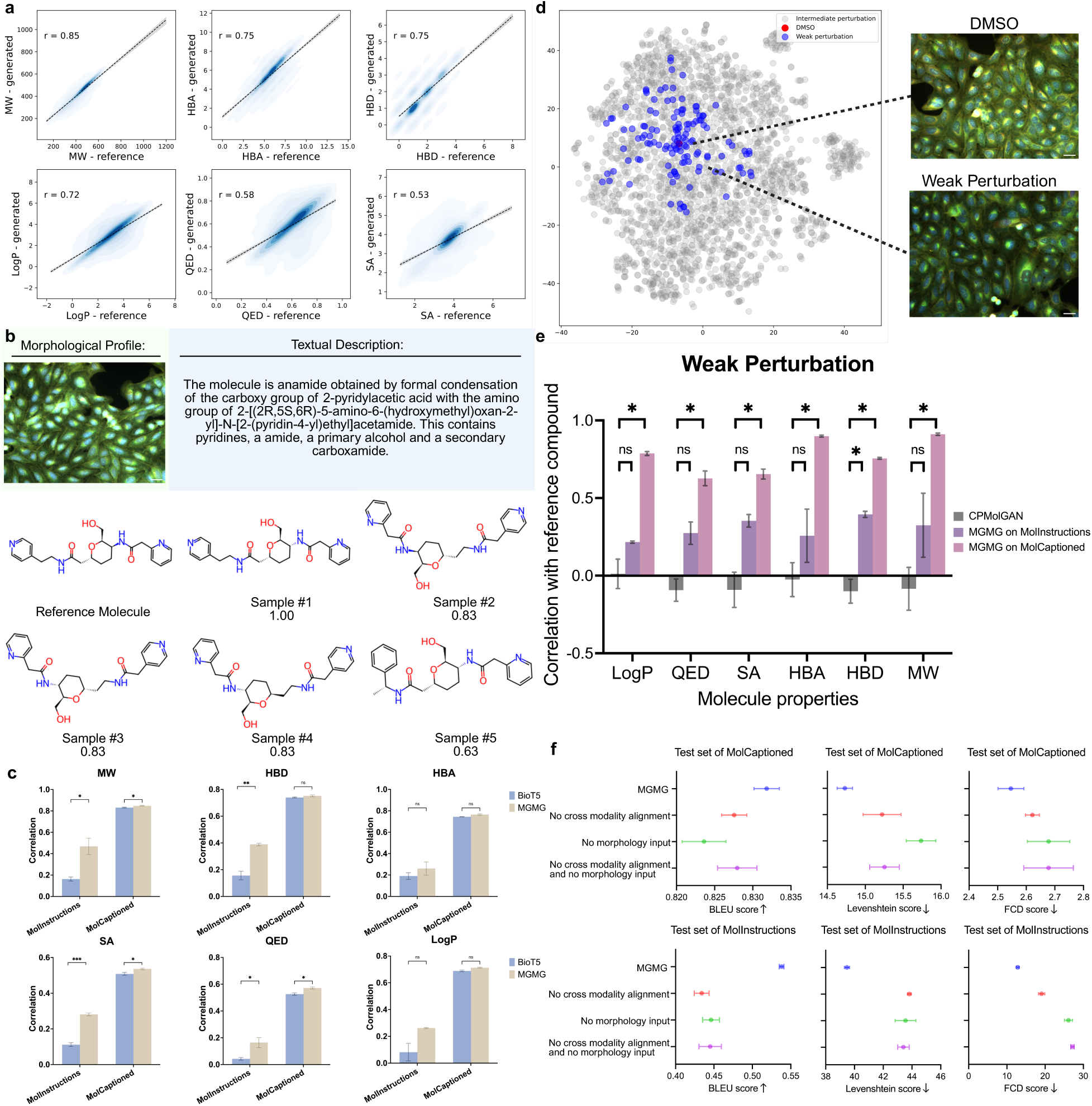
High Accuracy, Efficacy and Diversity in Molecule Generation by MGMG. **a** Properties of MGMG-generated molecules exhibit high correlation with their respective reference compounds. **b** An example demonstrating the diversity of MGMG-generated molecules: the top five molecules generated based on the textual description and morphological profile of a reference compound are shown, with the top-ranked molecule achieving the highest Tanimoto similarity to the reference compound. **c** Correlation between generated and reference compound properties across multiple chemical metrics. MGMG outperforms the unimodal baseline BioT5 on both datasets, with particularly pronounced improvements under limited-text conditions (Mol-Instructions-BBBC036v1). **d** t-SNE visualization of the molecules with weak morphological impact. **e** Correlation between generated and reference molecule properties with weak morphological impact. MGMG outperforms CPMolGAN across all evaluated chemical properties, with greater performance observed when detailed textual descriptions (MolCaptioned-BBBC036v1) were used. **f** Performance comparison on MolCaptioned-BBBC036v1 and Mol-Instructions-BBBC036v1 datasets with different ablation settings (y-axis). The x-axis shows the evaluation metrics: BLEU scores, Levenshtein scores, and FCD scores. MGMG achieves state-of-the-art (SOTA) performance in molecule generation across all metrics. All asterisks indicate statistical significance from student’s t-test: p < 0.05 (*), p < 0.01 (**), p < 0.001 (***), p < 0.0001 (****) and not significant (ns).

### MGMG generates molecules with diverse structures

Analysis across all generated outputs revealed that MGMG produced a higher percentage of unique molecules and exhibited greater Murcko scaffold diversity compared to baseline models (Table 1). These results suggest that MGMG generates structurally diverse molecules while preserving essential chemical properties. To further illustrate this capability, we analyzed the top five molecules generated from a single input consisting of a textual description and a morphological profile. The generated molecules were ranked based on their Morgan fingerprint similarity to the corresponding reference molecule (Fig. 2b). We observed that all five generated molecules shared a common structural core and functional groups with the reference molecule. Despite minor difference in stereochemistry, Sample-1 matched the reference molecule with a Tanimoto similarity score of 1.00, demonstrating MGMG’s ability to accurately recover reference molecular structures. Sample-2, 3, and 4 each achieved a Tanimoto similarity score of 0.83. These molecules, which are stereoisomers of one another, introduced a minor variation in the placement of the carbonyl group, while retaining the overall scaffold and functional motifs. Sample-5, with a lower Tanimoto similarity score of 0.63, incorporated more substantial structural changes, including the substitution of the pyridine ring with a phenyl ring and the introduction of a branched alkyl chain in contrast to the linear two-carbon chain in the reference, while still maintaining the general chemical framework observed in the higher-ranked samples. These findings indicate that MGMG maintains essential molecular scaffolds while incorporating structural diversity. Such variation is valuable for exploring novel analogs around a known active compound. Additional examples of top five generated molecules from various input conditions are provided in Supplementary Fig. 3.

### Cellular morphological information improves generated molecule properties

We next evaluated the contribution of incorporating morphological profiles to the properties of generated molecules. Molecules were generated under two input configurations: (1) combined molecular textual descriptions and morphological profiles, and (2) textual descriptions alone (Supplementary Table 1). To challenge the model in conditions where limited chemical structural information is available, we used molecular descriptions from Mol-Instructions-BBBC036v1^20^, a dataset containing under-informative, non-unique textual descriptions sourced from PubChem. Compared to the detailed captions in the MolCaptioned-BBBC036v1 dataset, these under-informative descriptions provide substantially less structural detail (Supplementary Fig. 4). This setup more accurately reflects real-world drug design scenarios, where prior knowledge of the specific structures and properties of the molecule to be designed is often limited.

To assess the contribution of morphological information, we compared the outputs of MGMG with BioT5, a unimodal baseline model trained on the same molecular descriptions but without morphological input. We evaluated the correlation between generated molecules and their corresponding reference compounds across multiple chemical properties, including QED, SA, MW, LogP, HBA, and HBD. MGMG consistently outperforms BioT5 across both datasets, with particularly pronounced improvements in the Mol-Instructions-BBBC036v1 setting, where textual guidance is minimal. Notably, MGMG achieved a 282.039% increase in QED correlation and a 152.084% increase in SA correlation over BioT5 on Mol-Instructions-BBBC036v1, compared to more modest gains of 8.600% and 5.507%, respectively, on MolCaptioned-BBBC036v1 (Fig. 2c). This is expected, as the inclusion of rich phenotypic information becomes especially valuable when textual inputs lack detailed cues, whereas in the MolCaptioned-BBBC036v1 dataset, textual descriptions already provide informative guidance for molecular generation. Similar trends were observed for MW, LogP, HBA, and HBD (Fig. 2c), further highlighting the benefits of integrating morphological input. These findings indicate that morphological information enhances the drug-likeness and the synthesizability properties of generated molecules, especially when the available textual descriptions lack specificity.

### Textual information enhances molecule generation when morphological signals are weak

While the addition of morphological information has been shown to improve molecule generation when textual descriptions are limited, it is equally important to assess whether incorporating textual descriptions can compensate in scenarios where morphological signals are weak. As previously discussed, weak perturbations often induce only subtle morphological changes, making it challenging to extract clear, actionable signals for molecule generation^15^. This limitation is especially relevant for models that rely exclusively on morphological input, such as CPMolGAN^15^. In such cases, their ability to generate meaningful compounds from weak morphological signals is limited^15^. In contrast, MGMG incorporates both morphological and textual descriptions, enabling it to compensate for weak phenotypic signals by leveraging textual information, thereby effectively enhancing the specificity and control in the molecule generation process. This dual-modality design allows MGMG to maintain generation performance even when morphological profiles are minimally informative.

To test this hypothesis, we quantified the strength of morphological perturbation by computing the Euclidean distance between each treatment profile and the DMSO control—a commonly used phenotypically inert reference. Compounds were then ranked in ascending order by their distance to DMSO, and the bottom 5% (n = 130) with the weakest morphological signals were selected for analysis (Fig. 2d). On this subset, MGMG substantially outperformed CPMolGAN, achieving significantly higher correlations between generated molecules and their reference counterparts across key chemical properties, such as QED, SA, MW, LogP, HBA, and HBD (Fig. 2e). These performance gains were especially pronounced when detailed textual descriptions (MolCaptioned-BBBC036v1) were provided, as expected, compared to the under-informative textual descriptions from the Mol-Instructions-BBBC036v1 dataset (Fig. 2e). These findings underscore the importance of integrating textual descriptions, especially when phenotypic signals are weak. By leveraging both modalities, MGMG demonstrates greater robustness than morphology-only approaches, offering a more reliable strategy for molecule generation regardless of perturbation strengths.

### Ablation studies for MGMG

To validate the design of MGMG, we conducted a series of ablation studies using both the MolCaptioned-BBBC036v1 and Mol-Instructions-BBBC036v1 datasets (Fig. 2f, Supplementary Table 1). Our first set of experiments examined the effect of incorporating morphological profiles on molecule generation performance. Using identical input text, we compared two input configurations: (1) text-only generation and (2) multimodal generation with both morphological profiles and textual descriptions. Across both datasets, the inclusion of morphological input consistently improves reconstruction performance. On MolCaptioned-BBBC036v1, multimodal generation resulted in a 0.998% improvement in BLEU score and a 6.397% reduction in Levenshtein distance compared to text-only generation. On Mol-Instructions-BBBC036v1, the gains were more substantial, with BLEU increasing by 20.583% and Levenshtein distance decreasing by 9.352% (Fig. 2f, Supplementary Table 1).

We further evaluated correlations between generated and reference molecules across key chemical properties. Drug-likeness improved with multimodal generation, as shown by the reductions in FCD: 2.546 vs. 2.678 on MolCaptioned-BBBC036v1 (−4.932%) and 12.835 vs. 26.112 on Mol-Instructions-BBBC036v1 (−50.846%) (Fig. 2f, Supplementary Table 1). MGMG-generated molecules also showed significantly higher QED correlations under the multimodal generation setting for both datasets. Notably, on Mol-Instructions-BBBC036v1, the incorporation of morphological profiles led to a 111.347% increase in QED correlation relative to the text-only baseline, while the gain on MolCaptioned-BBBC036v1 was more modest at 7.567% (Supplementary Table 1). A similar pattern was observed for SA, with correlation increases of 96.805% on Mol-Instructions-BBBC036v1 and 5.560% on MolCaptioned-BBBC036v1 (Supplementary Table 1). Consistent improvements were also observed across MW, LogP, HBA, and HBD, reinforcing the contribution of morphological profiles to molecule generation, particularly when textual descriptions are under-informative.

In a second ablation study, we investigated the role of modality alignment in multimodal generation. Specifically, we evaluated whether aligning the morphology and text encoders via contrastive learning would improve performance. The model with aligned encoders outperformed the version without alignment across both datasets. For instance, on MolCaptioned-BBBC036v1, the MGMG model with modality alignment yielded a 0.515% improvement in BLEU score, a 3.231% reduction in Levenshtein distance, and a 2.875% reduction in FCD (Fig. 2f). These improvements were more pronounced on Mol-Instructions-BBBC036v1, where BLEU score increased by 23.960%, Levenshtein distance decreased by 9.905%, and FCD dropped by 32.796% (Fig. 2f). These results underscore the importance of projecting both modalities into a unified latent space to maximize the effectiveness of multimodal conditioning in molecule generation.

### MGMG enables activator design with genetic perturbations

Given the success of MGMG in generating molecules guided by compound-perturbed morphological profiles, we next investigated whether the model could be applied to gene overexpression morphological profiles to design activators. We designed this case study not only to assess the model’s generalizability but also to challenge it to generate functionally relevant molecules without any prior knowledge of binding target or reference compound structure.

Specifically, we applied MGMG to nine genes: NFKB1, BRCA1, HSPA5, TP53, CREBBP, STAT1, STAT3, HIF1A, and NFKBIA, using gene overexpression image profiles from the BBBC037v1 morphological profiling dataset^35^. According to prior studies, three of the nine genes—NFKB1, BRCA1, and HSPA5— induce strong morphological changes, while the remaining genes show weaker differences compared to empty vector controls^15^. For the textual descriptions, we used the GPT-4.0^36^ to generate the functional descriptions of desired molecular effects for each gene, explicitly excluding any structural or scaffold-level hints, thereby mimicking a realistic early-stage drug design scenario where functional outcomes are defined but direct knowledge of reference compound is unavailable (Fig. 3a,b, Supplementary Table 2).

**Fig. 3:**
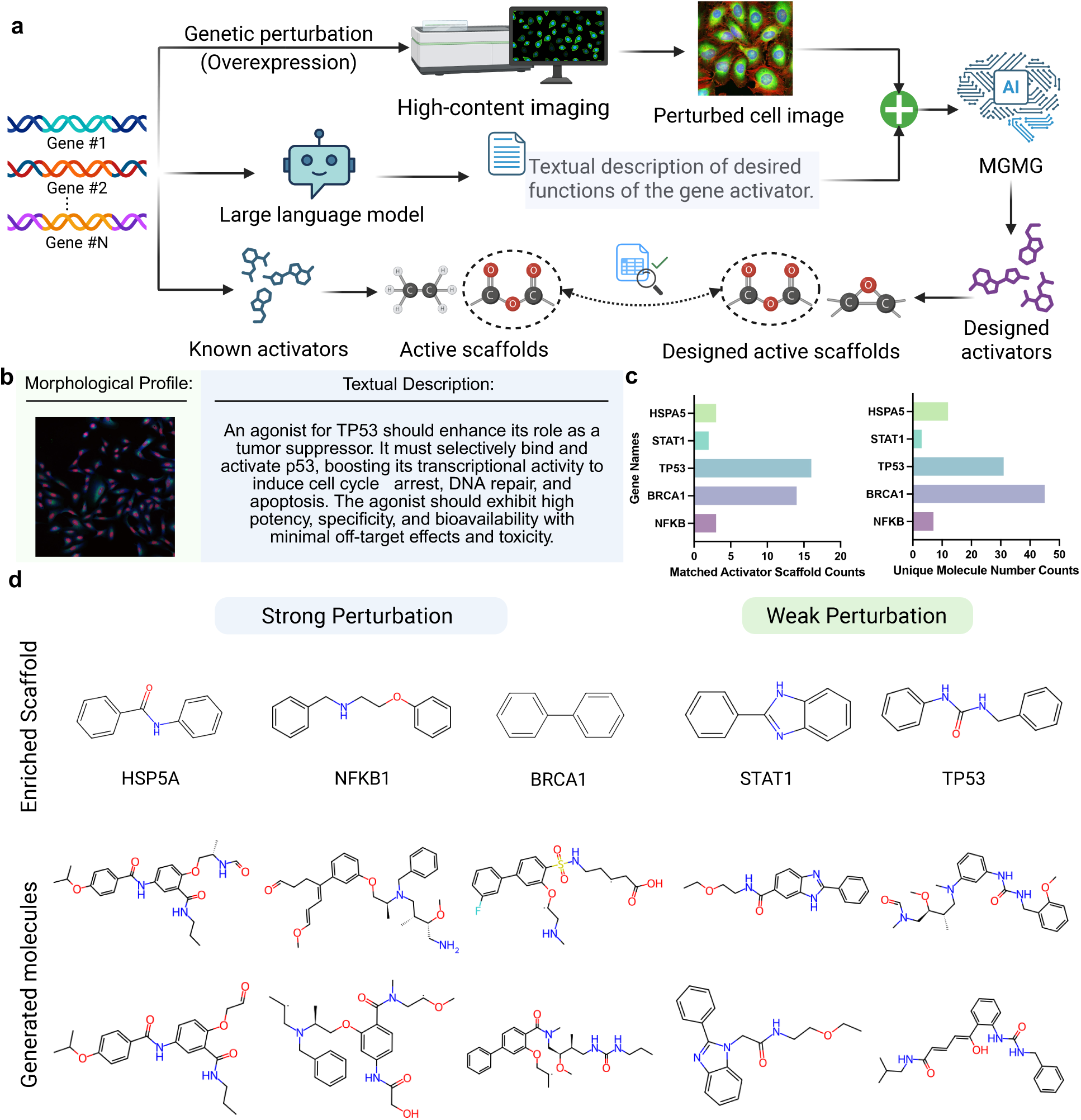
Molecules can be generated with gene overexpression morphological profiles and exhibit structural features resembling known activators. **a** Schematic overview of the activator design pipeline using MGMG. **b** An example input pair showing TP53 gene overexpression image and textual description detailing the desired properties for the activators to be designed. **c** Number of enriched scaffolds as in ExCAPE activators and number of corresponding molecules generated per gene. Representative generated molecules from genes with **d** strong and **e** weak morphological impacts, respectively.

To evaluate the quality and relevance of the generated molecules, we compared their scaffolds to known activators retrieved from the ExCAPE dataset^37^, assuming that gene overexpression can phenocopy activator activity (Fig. 3a). All known activators were confirmed to be absent from the MGMG training set (BBBC036v1), ensuring that scaffold recovery was not due to data leakage or memorization. Among the nine genes tested, MGMG successfully recovered ExCAPE activator scaffolds for five genes, outperforming CPMolGAN, which recovered activators for two^15^ (Fig. 3c). Notably, MGMG identified enriched scaffolds not only for genes with strong morphological signals such as NFKB1, BRCA1, and HSPA5 (Fig. 3d), but also for genes with weaker morphological impact, such as STAT1 and TP53 (Fig. 3e). For STAT1, two generated scaffolds matched potent activators in the ExCAPE dataset; for TP53, 16 generated scaffolds matched known activators (Supplementary Table 2).

These findings demonstrate that MGMG can be applied beyond chemical perturbations to genetic perturbation profiles and design molecules that resemble known activators even in the absence of their structural information. This capability underscores MGMG’s unique target-agnostic potential, enabling molecule generation based solely on phenotypic outcomes and functional goals. This target-agnostic approach opens new possibilities for molecule design, particularly in cases where structure-activity relationships are unknown or where conventional target-based approaches are limited.

### Molecular docking of MGMG-generated molecules to potential protein targets

Although MGMG is a target-agnostic molecule design approach, here we investigated whether the designed molecules could bind to potential targets when the reference compounds’ targets are available. For this case study, we docked the molecules generated by MGMG to the protein targets of the reference compounds, where the corresponding protein structures were fetched from the Protein Data Bank (PDB), and then we compared the docking results to those of the reference molecules. To ensure adequate structural diversity, the top five designed molecules were selected for docking. Docking scores were calculated by Glide SP^38^, based on the co-crystallized or docking-generated poses.

In the first case (Fig. 4a), the reference molecule BRD-K85883481-001-02-6 was docked to its receptor, human liver 5β-reductase (AKR1D1) (PDB ID: 3CMF), yielding a docking score of -8.81 kcal/mol. The generated Sample-1 molecule retained the same steroid-like core structure as the reference molecule and only replaced the ketone on the 11-position by a hydroxyl. In contrast, the Sample-2 molecule demonstrated greater structural divergence, replacing the ketone on the 11-position with a hydroxyl group and removing the alpha hydroxyl group on the 17-position. Despite these differences, both the Sample-1 and Sample-2 molecules successfully retained the reference molecule’s key interactions with the target protein, including hydrogen bonds to SER169 and TYR58. These preserved interactions stabilized their binding within the protein pocket, resulting in docking scores of -8.46 kcal/mol for the Sample-1 molecule and -8.50 kcal/mol for the Sample-2 molecule (Fig. 4a). These results demonstrate MGMG’s capacity to generate structurally diverse molecules with similarly promising docking scores, highlighting its potential to explore novel chemical space.

**Fig. 4:**
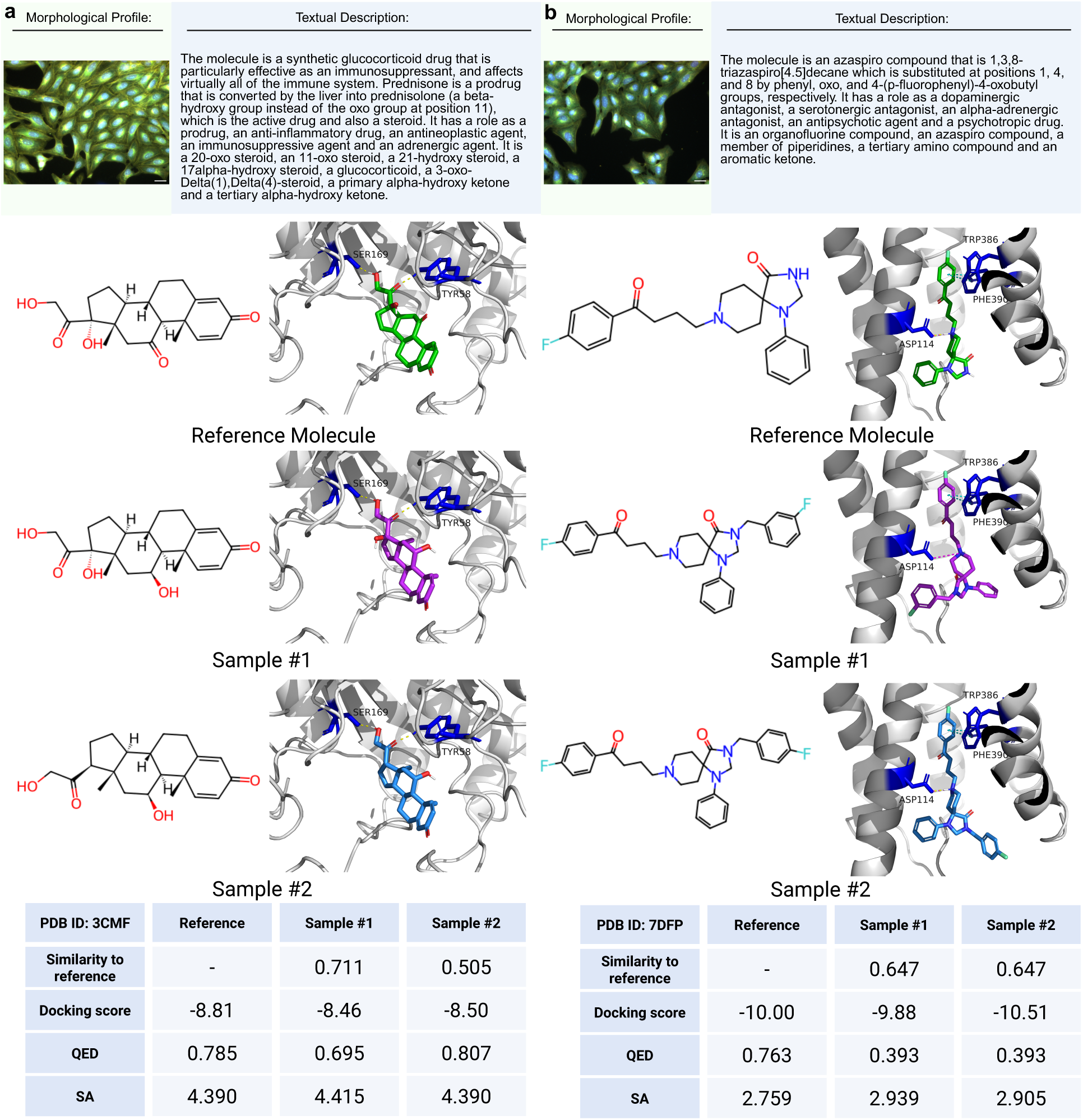
Molecular docking of MGMG-generated molecules to potential protein targets. The top panel presents the input textual descriptions and cell morphology profiles of reference molecules targeting: **a** human liver 5β-reductase (PDB ID: 3CMF) and **b** human dopamine D2 receptor (PDB ID: 7DFP). The middle panel displays molecular structures and three-dimensional docking results of the ligands in their best-scored binding poses within the respective binding pockets. Proteins are shown in a grey cartoon representation, while ligands are shown as stick models. Carbon atoms are colored green for reference molecules and pink or light blue for MGMG-generated molecules. Oxygen and nitrogen atoms are colored red and blue, respectively. The target protein residues involved in interactions are highlighted in dark blue. Interaction types are represented as follows: hydrogen bonds (yellow dashed lines), salt bridges (magenta dashed lines), and Pi-Pi interactions (cyan dashed lines). The bottom panel quantifies molecular similarity and drug-like properties, comparing MGMG-generated molecules to the reference molecules based on Tanimoto similarity, docking scores, QED, and SA.

In another case (Fig. 4b), the reference molecule BRD-K55468218-001-04-8 was docked to the human dopamine D2 receptor (PDB ID: 7DFP), with a docking score of -10.00 kcal/mol. Compared to the reference molecule, Sample-1 and Sample-2 molecules both included an additional fluorobenzene ring, differing only in the fluorination position. This structural extension enabled the generated compounds to reach subpockets not accessed by the reference molecule. Despite these structural differences, both designs preserved the reference compound’s key interactions with the target protein, including the salt bridge and hydrogen bond between the positively charged nitrogen and ASP114, as well as the Pi-Pi stacking interactions with TRP386 and PHE390. These interactions resulted in docking scores of -9.88 kcal/mol and -10.51 kcal/mol, respectively. This case highlights MGMG’s potential for scaffold expansion to improve target engagement and explore new binding regions.

Additional cases are presented in Supplementary Fig. 5, further demonstrating the capability of MGMG to produce diverse molecules that not only maintain comparable binding affinities to the reference molecules but also introduce meaningful structural variations. Collectively, these findings show that molecules generated by MGMG, despite lacking explicit knowledge of the target protein, can retain the critical interactions with key target residues, thereby ensuring stable binding. At the same time, the structural variations introduced by MGMG allow the molecules to explore novel chemical space and previously untapped regions of the binding pocket, supporting the dual goals of structural novelty and optimization in ligand design.

## Discussion and conclusion

Developing novel molecules with favorable chemical properties and bioactivity remains a longstanding challenge in drug discovery. Most existing in silico molecule design approaches follow the principles of target-based drug discovery, relying on well-defined disease targets and therefore offering limited applicability for diseases without validated molecular targets. In this study, we present MGMG, a phenotypic drug discovery-oriented framework that integrates morphological information from compound treatment with molecular textual descriptions to enable *de novo* molecule design without the need for target information. A comprehensive benchmark study demonstrates that our approach outperforms state-of-the-art models that rely solely on either morphological information or textual descriptions. Detailed ablation study further confirms that both the incorporation of morphology data and the multimodal alignment via contrastive learning contribute to the model’s improved performance. To illustrate the practical applicability of MGMG beyond benchmark evaluations, we present a case study in activator design, demonstrating how this framework not only can be applied from chemical to genetic perturbations but also enables target-agnostic molecule generation without requiring any structural information from reference compounds, relying solely on morphological and functional cues. In addition, we show that, despite the absence of explicit target structural knowledge, MGMG-generated molecules can preserve key interactions with target residues while introducing structural variations to explore novel chemical space.

In this study, we propose the integration of cell morphology and molecular textual description modalities for target-agonistic molecule generation. These two distinct modalities address different aspects of the molecule design task and effectively complement each other. Cell morphology data, such as from Cell Painting assays, provide a comprehensive overview of cellular responses to compound treatment, effectively reflecting compound bioactivity. Large public datasets, such as the JUMP-CP consortium^39^, have been released to provide cellular morphological profiles from both chemical and genetic perturbations^12^. Textual descriptions provide a direct and often interpretable characterization of molecule structure. Recent datasets derived from PubChem and ChEMBL^20, 40^ have significantly expanded the availability of text-molecule pairs; however, these descriptions typically offer limited insight into biological functions of the molecules. One of the key findings of our study is the complementary role that morphological and textual contexts play in molecule generation, particularly under limited-information scenarios. In real-world settings, it is common for one source of inputs to be under-informative—either from non-specific textual descriptions due to limited prior knowledge of the reference compound, or from weak phenotypic perturbations that produce minimal morphological changes. Our experiments show that morphological profiles significantly enhance generation performance when textual description lacks structural details, and that textual descriptions can meaningfully compensate for low-information morphological profiles from phenotypically inert compounds. This mutual compensatory effect addresses another key practical challenge in molecule generation, where consistent and informative guidance is lacking from either context alone. By integrating both morphological and textual information, MGMG offers a robust, target-agonistic strategy for designing bioactive molecules.

Although MGMG has demonstrated strong potential, it still faces several limitations that warrant further investigation. First, the current implementation relies on morphological profiles extracted using CellProfiler^22^, which requires additional software setup and manual parameter tuning. Moreover, these profiles are limited to a predefined set of hand-engineered features, which may not fully capture the rich and nuanced information embedded in the raw cell images. Leveraging end-to-end deep learning approaches, such as convolutional neural networks^41^ or vision transformers^12, 42^ applied directly to Cell Painting images, could enable the model to learn more unbiased, expressive and biologically relevant representations, potentially enhancing its generalizability and performance. Another limitation of MGMG lies in its reliance on morphological profiles derived from the single cell line U2OS, which may limit the model’s ability to generalize across diverse cellular contexts. Incorporating morphological data from multiple cell types in future studies would allow the model to better capture context-dependent phenotypic responses to compounds, leading to a more comprehensive and biologically informed molecular design framework^39^.

Conventional *de novo* molecular design methods based on TDD often fall short in complex, target-agnostic settings, particularly for diseases without validated targets or known reference compounds^10^. Through leveraging both morphological profiles and textual descriptions, our PDD-oriented framework addresses this gap by capturing holistic biological responses and enhancing molecule design in a target-independent manner, opening new directions for future development. With its demonstrated effectiveness, MGMG holds strong promise for driving target-agnostic molecule generation and expanding the reach of phenotypic-based approaches.

## Author Contributions

Y.L. conceived and supervised the research. Y.L. and Q.T. designed and implemented the methodology. Q.T. and D.D. implemented the baselines. X.Y., C.L., and H.L. contributed to discussions on experimental design. P.R. and C.L. refined the molecular textual descriptions. Y.L., Q.T. and G.S. wrote the paper. All authors reviewed and approved the manuscript.

## Conflicts of Interests

The authors do not have conflicts to declare.

## Data Availability

All data supporting the findings of this study are available on Zenodo at https://zenodo.org/records/16296425. The dataset includes the trained model checkpoint, test set inputs of aligned morphological profiles and molecular text descriptions, and input materials for both the activator design and docking studies. The code used to generate the results presented in this study is provided as a supplementary file.

## Acknowledgements

This work was supported in part by the University of Florida (UF Startup Fund and UF Research AI Award to Y. L., UF Blue Future Medicine Initiative to H. L.), the Florida Department of Health (Florida Cancer Innovation Fund grant #MOABK), National Institutes of Health (R21EB037868 to Y. L. and X. Y. and RM1GM145426 to H. L.), the Debbie and Sylvia DeSantis Chair Professorship (to H. L.), the Bodor Professorship Fund (to C. L.), and the National Science Foundation (NSF South Carolina EPSCoR 25-GA04). We acknowledge UFIT Research Computing for providing computational resources. Figures 1, 2, 3, 4 were created with BioRender.com.

## Notes

### Competing Interest Statement

The authors have declared no competing interest.

### Summary of Updates

Supplementary Information, including notes, figures, and tables, has been uploaded.

