## Supplementary Information for "MGMG: Cell Morphology-Guided Molecule Generation for Drug Discovery"

* Correspondence: Yanjun Li

### SUPPLEMENTARY NOTES

#### Dataset construction

##### *Mol-Instructions-BBBC036v1 and MolCaptioned-BBBC036v1 datasets*

In this study, we curated two tri-modal datasets, which aligned the molecular structure (represented as SELFIES), the corresponding textual description, and the induced cellular morphological profile under compound treatment. Wherein, the MolCaptioned-BBBC036v1 dataset was used for both model training and evaluation, while the Mol-Instructions-BBBC036v1 dataset was used exclusively for evaluation. We started from the BBBC036v1 dataset from the Broad Bioimage Benchmark Collection, which includes morphological profiles of 30,616 compounds treated on the U2OS cells in replicates and imaged after Cell Painting assay[1]. The morphological profiles were processed with the Pycytominer pipeline for normalization, feature selection (reduction) and replicated well consensus[2]. From this pipeline, we reduced feature dimension to 753. To acquire the textual descriptions for these compounds, their canonical SMILES representations were utilized to match corresponding entries in the Mol-Instructions[3] datasets. A total of 25,070 compounds were identified to have both the morphological profiles from BBBC036v1 and textual descriptions from PubChem, forming the Mol-Instructions-BBBC036v1 dataset[3]. To improve the description quality, we used pre-trained molecular captioning model BioT5 to generate molecule captions for these 25,070 compounds, resulting in the creation of MolCaptioned-BBC036v1 dataset. We partitioned this tri-modal dataset into training, validation and testing sets with an 8:1:1 ratio.

#### MGMG model and training details

##### *MGMG encoder*

The encoder of MGMG follows the transformer-based architecture of BioT5[4], which is built on the T5-v1.1-base[5] configuration. The encoder consists of 12 transformer layers, each comprising a multi-head self-attention mechanism and a feed-forward network. Each attention block includes 12 attention heads, with key and value projection dimensions of 64, and the model-wide hidden state dimension is set to 768. The feed-forward network within each block consists of a dense layer of size 3072 followed by a ReLU activation and a second dense layer projecting back to 768 dimensions. A dropout probability of 0.1 is applied throughout for regularization. The output of the encoder is passed to the decoder for autoregressive molecule generation.

*Morphological prefix and early fusion strategy*

To integrate phenotypic information, the cellular morphological profile extracted from CellProfiler[6] is projected into a 768-dimensional embedding space. The morphological embedding is then **prepended to the input text sequence** as the morphological prefix, forming a unified input to the MGMG encoder. This **early fusion strategy** allows the Transformer to jointly attend over both morphological and textual representations from the very first layer of processing.

*MGMG decoder*

The decoder of MGMG is structurally consistent with the encoder and also follows the T5-v1.1-base[5] configuration. It comprises 12 decoder layers, each with multi-head self-attention, cross-attention (attending to encoder outputs), and a feed-forward network. Dropout of 0.1 is applied uniformly. The decoder takes as input the embedded representation of SELFIES tokens and performs autoregressive decoding. The final decoder output is projected back to the SELFIES vocabulary space using a linear layer and softmax activation function, producing one token at a time during inference. This setup allows MGMG to generate chemically valid molecules conditioned on both textual description and morphological context.

*Modality alignment with contrastive learning*

In the first stage of training, MGMG learned to align the two input modalities—textual descriptions and cellular morphological profiles—by projecting them into a shared latent space. This was achieved using a contrastive learning approach by optimizing the InfoNCE[7] loss function.

Given a batch of size N, we computed the pairwise similarity scores between text and morphology embeddings. Let $S_{ii}$ denote the similarity between a correctly aligned (positive) text–morphology pair, and $S_{ij}$ represent the similarity between mismatched (negative) pairs. A temperature hyperparameter $\tau$ controlled the sharpness of the similarity distribution. The contrastive loss was defined as:

$$L_{\text{contrastive}} = -\frac{1}{N}\sum_{i=1}^{N}\log\left(\frac{\exp(s_{ii}/\tau)}{\sum_{j\neq i}\exp\left(s_{ij}/\tau\right)}\right)$$

We used a batch size of 48 and the temperature hyperparameter $\tau$ was set as 0.1. The weights of the morphology encoder from this stage were subsequently used in the molecule generation stage, where both modalities were jointly processed in an autoregressive decoding framework.

##### *Training for molecule generation*

To train the MGMG model for molecule generation, we used the AdamW optimizer and applied a cosine annealing learning rate scheduler, starting at a base learning rate of $1 \times 10^{-3}$ and decaying to a minimum of $1 \times 10^{-5}$. The batch size was 48. To reduce overfitting and improve parameter efficiency, Low-Rank Adaptation (LoRA) was applied during training with a rank of 128, alpha value of 128, and a dropout rate of 0.1. This reduced the number of trainable parameters to 14 million—just 5.3% of the full model's 267 million parameters. Training was performed on one NVIDIA A100 (80G) GPU. The MGMG model was trained for ~90 epochs, totaling ~51 hours of wall clock time. Training and evaluation were repeated using three different random seeds to ensure reproducibility.

#### Baseline models

We benchmarked MGMG against a range of baseline models, including MolT5, BioT5, and CPMolGAN. Each model was trained on the MolCaptioned–BBBC036v1 dataset using its own default or published hyperparameter configurations. To ensure reproducibility, training and evaluation for each model were repeated across three independent random seeds.

- CPMolGAN[8] is the first model to use cell image features that capture compound-treated cell morphology as the input for molecule generation. In this framework, morphological profiles derived from the Cell Painting assay are used to condition the GAN, guiding the generation of molecules toward desired phenotypic effects[8].
- MolT5[9] is built upon the T5 architecture for joint modeling of natural language and molecular representations. It is pretrained on the "Colossal Clean Crawled Corpus" (C4)[5] dataset for the text modality and the ZINC dataset for molecule modality, enabling tasks such as molecule captioning and text-based molecule generation.
- BioT5[4] is a T5-based model that learns from both molecular structures and biomedical text. It is pretrained on **339K molecule–text pairs from PubChem**, where each molecule is represented as a **SELFIES** string paired with its corresponding textual description[4]. This ensures chemically valid generation while enabling the model to capture meaningful associations between molecular structures and biological literature.

#### Evaluation Metrics

To evaluate the performance of MGMG in molecule generation, we used the following metrics:

##### (1) Reconstruction metrics:

- **BLEU score.** BLEU (Bilingual Evaluation Understudy) is a precision-based metric that quantifies the similarity between the generated and reference molecular sequences. It measures the overlap of n-grams while applying a brevity penalty to discourage excessively short outputs[10]. A higher BLEU score indicates that the generated sequences more accurately reconstruct the reference molecular sequences.
- **Levenshtein distance.** The Levenshtein distance measures the minimum number of single-character edits (insertions, deletions, and substitutions) required to transform the

generated molecular sequence into the reference molecular sequence[11]. A lower Levenshtein distance indicates a closer sequential similarity to the reference molecule.

##### (2) Synthesizability:

The **Synthetic Accessibility (SA) score** quantifies how easily a given molecule can be synthesized, with lower scores indicating greater synthetic feasibility[12]. Following the approach of Zapata et al.[8], we applied a threshold of 4.5 to report the percentage of compounds considered synthetically accessible.

##### (3) Drug-likeness:

- **Fréchet ChemNet Distance (FCD).** FCD assesses the structural similarity between generated and real molecules by comparing their latent space representations obtained from a pretrained ChemNet model. It quantifies how closely the distribution of generated molecules resembles that of real, bioactive compounds. Specifically, for two sets of molecules—generated set $G$ and reference set $R$—the FCD is computed as the Fréchet distance between their multivariate Gaussian embeddings:

$$\mathrm{FCD}(G,R) = |\mu_G - \mu_R|^2 + \mathrm{Tr}\left[\Sigma_G + \Sigma_R - 2(\Sigma_G \Sigma_R)^{1/2}\right]$$

  where $\mu_G$, $\mu_R$ are the mean vectors and $\Sigma_G$, $\Sigma_R$ are the covariance matrices of the latent activations for the generated and reference molecules, respectively. A lower FCD indicates that the generated molecules are more structurally and chemically similar to real compounds[13]. The PyTorch implementation provided by Insilico Medicine (https://github.com/insilicomedicine/fcd_torch) was used to compute this metric.

- **Quantitative Estimate of Drug-likeness (QED):** The QED score measures the drug-likeness of a molecule based on multiple physicochemical properties. Higher QED scores indicate stronger alignment with known drug-like characteristics[14]. In this work, we followed prior conventions[8] and applied a threshold of 0.5 to quantify the percentage of generated molecules with favorable drug-like properties.

##### (4) Physicochemical properties:

We computed Molecular Weight (MW), Lipophilicity (LogP), Hydrogen Bond Acceptors (HBA) and Hydrogen Bond Donors (HBD) using RDKit[15].

##### (5) Diversity:

To assess the structural diversity of the generated molecules, we computed **Murcko scaffolds**[16] using RDKit, which extracts the core molecular structure by removing side chains while retaining ring systems and linkers. Diversity is defined as the proportion of unique scaffolds among all valid generated molecules, reflecting the model's ability to explore diverse chemical backbones rather than repeatedly generating similar structures[17].

##### (6) Validity:

Validity is defined as the proportion of generated molecular strings, e.g., SMILES, that can be successfully parsed into chemically valid molecules using RDKit. A molecule is considered valid if it satisfies fundamental chemical rules, including atom valency and bond configurations, particularly within aromatic systems. This metric assesses the model's ability to generate syntactically correct and chemically feasible structures[17].

##### (7) Uniqueness:

Uniqueness measures the proportion of unique molecules among all the valid generated molecules. This metric ensures that the model does not collapse into generating redundant or repetitive structures and reflects its capacity for diverse molecule generation[17].

#### Molecule generation for activator design task

For the activator design study, we used gene overexpression morphological profiles from Zapata et al.'s work[8]. These profiles were originally derived from the BBBC037v1 dataset from the Broad Bioimage Benchmark Collection, which contains Cell Painting images of U2OS cells transfected to overexpress 220 genes[18]. We focused on nine genes with known matched activators in the ExCAPE activators[19] database. All profiles were processed using the normalization and aggregation strategy described in Zapata et al.'s work[8]. Phenyl rings were excluded because they are ubiquitous hydrophobic substituents and therefore provide limited information for scaffold enrichment analysis. Regarding the textual descriptions, we used GPT-4.0 to generate simple functional annotations for the desired activators, avoiding explicit structural cues. These input textual descriptions are provided in Supplementary Table 2.

For each gene, 2,500 valid molecules were generated using consensus morphological profiles. The process followed three steps to ensure chemical and semantic diversity: (1) 50 molecules were generated based on the GPT-4.0-provided descriptions and the gene's morphological profile; (2) these molecules were then captioned using BioT5; and (3) each caption was combined again with the same morphology profile to generate 50 additional molecules. All generated molecules were filtered using physicochemical criteria following Zapata *et al.*[8], including: $-2 < \text{LogP} < 7$, HBA + HBD < 10, topological polar surface area (TPSA) < 150, rotatable bonds < 150, and no SureChEMBL alerts.

#### Molecular Docking

Glide docking[20] was performed to assess interactions between the target protein and MGMG-designed molecules as well as reference molecules. Protein structures were prepared using the Schrödinger Protein Preparation Workflow at pH 7.4. Protonation states of ionizable amino acids were predicted using PROPKA, and those of ligands were determined using Epik. Waters beyond 5Å from ligands were removed, and the structure was minimized with the OPLS4 force field, converging heavy atoms to a root-mean-square deviation (RMSD) of 0.3Å . The receptor grid box was generated based on the binding site of the reference molecule with default Glide Grid Generation settings. MGMG-designed molecules were prepared for docking using Schrödinger LigPrep with the OPLS4 force field. Preparation included converting 2D to 3D structures, adding hydrogens, computing partial charges, and optimizing the structures. The docking results were reported as GlideScores. All calculations were conducted using Maestro (version 14.2.118, Schrödinger).

### SUPPLEMENTARY TABLES & FIGURES

| Property | Details | MolCaptioned-BBBC036v1 | | Mol-Instructions-BBBC036v1 | |
|---|---|---|---|---|---|
| | | With morphological features | Without morphological features | With morphological features | Without morphological features |
| Reconstruction metrics | BLEU score↑ | **0.832±0.003** | 0.824±0.005 | 0.538±0.005 | 0.446±0.019 |
| | Levenshtein score↓ | **14.730±0.176** | 15.737±0.338 | 39.476±0.314 | 43.549±1.248 |
| Physiochemical properties | MW | 430.712±0.560 | 431.648±2.275 | 391.069±7.346 | 367.677±6.628 |
| | MW $R^2$ to reference↑ | **0.848±0.005** | 0.834±0.010 | 0.469±0.133 | 0.260±0.050 |
| | LogP | 2.956±0.035 | 2.998±0.015 | 2.895±0.109 | 2.974±0.163 |
| | LogP $R^2$ to reference↑ | **0.713±0.007** | 0.685±0.018 | 0.262±0.007 | 0.183±0.084 |
| | HBA | 5.466±0.042 | 5.463±0.022 | 4.766±0.100 | 4.446±0.287 |
| | HBA $R^2$ to reference↑ | **0.765±0.012** | 0.750±0.010 | 0.260±0.107 | 0.253±0.026 |
| | HBD | 1.777±0.012 | 1.776±0.020 | 1.498±0.022 | 1.418±0.174 |
| | HBD $R^2$ to reference↑ | **0.752±0.013** | 0.736±0.006 | 0.388±0.015 | 0.236±0.110 |
| Drug-likeness | ChEMBL FCD↓ | **2.546±0.078** | 2.678±0.128 | 12.835±0.642 | 26.112±1.814 |
| | QED↑ | 0.589±0.003 | 0.590±0.001 | 0.620±0.015 | **0.636±0.035** |
| | %QED>0.5↑ | 0.714±0.009 | 0.714±0.005 | 0.763±0.037 | **0.783±0.025** |
| | $R^2$ to reference↑ | **0.572±0.013** | 0.531±0.022 | 0.164±0.064 | 0.078±0.066 |
| Synthesizability | SA score↓ | 3.695±0.014 | 3.731±0.006 | 3.661±0.028 | **3.494±0.066** |
| | %SA score<4.5↑ | **0.885±0.007** | 0.874±0.005 | 0.850±0.028 | 0.877±0.069 |
| | $R^2$ to reference↑ | **0.536±0.008** | 0.508±0.031 | 0.282±0.014 | 0.143±0.054 |
| Diversity | %Unique Murcko Scaffold↑ | **0.748±0.003** | 0.741±0.002 | 0.441±0.027 | 0.131±0.002 |
| Validity | %Valid↑ | **1.00±0.00** | **1.00±0.00** | **1.00±0.00** | **1.00±0.00** |
| Uniqueness | %Unique↑ | **0.980±0.001** | 0.975±0.002 | 0.613±0.018 | 0.155±0.0002 |

**Supplementary Table 1: Top 1 generated molecules by MGMG with or without morphology information on MolCaptioned-BBBC036v1 and Mol-Instructions-BBBC036v1 dataset.**

| Genes | Textual Description of Agonist | Valid Hit # matched to ExCAPE | Enriched Agonist Scaffold (in SMILES) | Uni mol # with scaffold |
|---|---|---|---|---|
| HSPA5 | An agonist for HSPA5 should enhance its role as a molecular chaperone, aiding in protein folding and ER stress response. It must selectively bind and activate HSPA5, improving its ability to stabilize nascent proteins and prevent misfolding. The agonist should be potent, specific, and have minimal toxicity. | 3 | c1ccc(COc2ccccc2)cc1 | 2 |
| | | | c1ccncc1 | 8 |
| | | | O=C(Nc1ccccc1)c1ccccc1 | 2 |
| STAT1 | An agonist for STAT1 should enhance its role in mediating cytokine signaling. It must selectively bind and activate STAT1, boosting its phosphorylation, dimerization, and transcriptional activity to promote expression of interferon-stimulated genes. The agonist should exhibit high potency, specificity, and minimal off-target effects. | 2 | c1ccc2ccccc2c1 | 1 |
| | | | c1ccc(-c2nc3ccccc3[nH]2)cc1 | 2 |
| TP53 | An agonist for TP53 should enhance its role as a tumor suppressor. It must selectively bind and activate p53, boosting its transcriptional activity to induce cell cycle arrest, DNA repair, and apoptosis. The agonist should exhibit high potency, specificity, and bioavailability with minimal off-target effects and toxicity. | 16 | O=C(NCc1ccccc1)Nc1ccccc1 | 2 |
| | | | O=C(NC1CCCCC1)c1ccccc1 | 1 |
| | | | c1ccc(-c2ccco2)cc1 | 4 |
| | | | c1ccc(Oc2ccccc2)cc1 | 1 |
| | | | c1ccc(CCNc2ccccc2)cc1 | 1 |
| | | | c1ccc(N2CCNCC2)cc1 | 2 |
| | | | c1ccc(-c2ccccc2)cc1 | 4 |
| | | | O=C(NCCOc1ccccc1)c1ccccc1 | 1 |
| | | | O=C(Cc1ccccc1)NCc1ccccc1 | 2 |
| | | | c1ccncc1 | 7 |
| | | | O=C(Nc1ccccc1)c1nc2ccccc2[nH]c1=O | 1 |
| | | | c1ccc2nc3ccccc3cc2c1 | 1 |
| | | | O=c1ccc2ccccc2o1 | 1 |
| | | | c1ccc(OCCOc2ccccc2)cc1 | 1 |
| | | | O=C(Nc1ccccc1)Nc1ccccc1 | 1 |
| | | | O=C(Nc1ccccc1)C1CC1 | 1 |
| BRCA1 | An agonist for the BRCA1 gene should enhance its DNA repair capabilities, stabilize genomic integrity, and reduce mutation rates. The agonist must selectively bind to and activate key functional domains of the BRCA1 protein, such as the RING and BRCT domains, boosting its interaction with other DNA repair proteins. It should exhibit high potency, specificity, and bioavailability while minimizing off-target effects and toxicity. The ideal agonist would promote efficient DNA damage repair, decrease cancer risk, and ensure cellular stability without disrupting other cellular processes. | 14 | O=C(Nc1ccccc1)c1ccccc1 | 1 |
| | | | O=C(Nc1ccccc1)c1ccon1 | 5 |
| | | | c1ccc(-c2ccccc2)cc1 | 3 |
| | | | c1ccncc1 | 7 |
| | | | O=C(Nc1ccccc1)c1ccccn1 | 1 |
| | | | c1ccc(COc2ccccc2)cc1 | 1 |
| | | | c1ccc(-c2cccnc2)cc1 | 20 |
| | | | O=C(NCCOc1ccccc1)c1ccccc1 | 1 |
| | | | O=C(NCCC(=O)N1CCCCC1)c1ccccc1 | 1 |
| | | | O=C(COc1ccccc1)c1ccccc1 | 1 |
| | | | O=C(NCCc1ccccc1)c1ccccc1 | 1 |
| | | | O=C(CNC(=O)c1ccccc1)c1ccccc1 | 1 |
| | | | c1ccc2c(c1)OCCO2 | 1 |
| | | | c1ccc(CN2CCCCC2)cc1 | 1 |
| NFKB1 | An agonist for NFKB1 should enhance its role in regulating immune response and inflammation. It must selectively bind and activate p50 subunit, boosting NFKB transcriptional activity to promote expression of immune and inflammatory genes. High potency, specificity, and bioavailability with minimal off-target effects are crucial. | 3 | c1ccc(Oc2ccccc2)cc1 | 1 |
| | | | c1ccc(CNCCOc2ccccc2)cc1 | 2 |
| | | | c1ccncc1 | 4 |

**Supplementary Table 2: Textual description inputs and matched scaffold hits in the activator design case study.**

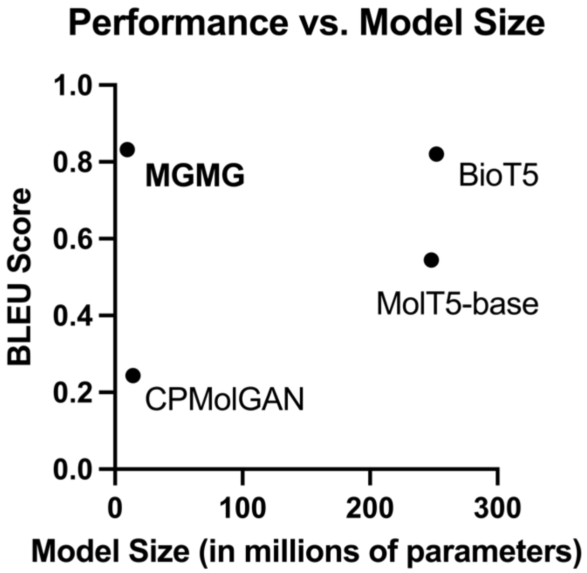


**Supplementary Fig. 1: Comparison of MGMG and unimodal baselines on trainable parameter counts and molecule reconstruction performance as measured by BLEU score.**

| Textual Description | Morphological Profile | Output | | | | |
|---|---|---|---|---|---|---|
| The molecule contains carboxamides obtained by the formal condensation of the carboxy group of el-Methyl (1R,3R,4aS,9aR)-3,4,4a,9a-tetrahydro-1-(hydroxymethyl)-6-[(2-methoxyacetyl)amino]-1H-pyrano[3,4-b]benzofuran-3-acetate with the amino group piperonylamine. This contains a carboxamide and benzodioxoles. It derives from a (+)-alpha-amino-L-tyrosine. | 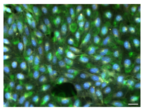 | BRD-K01248935-001-01-7 Reference Molecule | MGMG 0.90 | BioT5 0.68 | MolT5 0.22 | CPMolGAN 0.15 |
| The molecule is an azabicycloalkane that is the 4-methoxybenzoate ester of a carbamic acid. A cyclooxygenase 2 inhibitor, it is used in the treatment of chronic obstructive pulmonary disease (COPD) and gout. It has a role as a cyclooxygenase 2 inhibitor, a non-narcotic analgesic, a gout suppressant and a xenobiotic. It is an aromatic ether, a carbamate ester, a member of triazoles, and a tertiary carboxamide. | 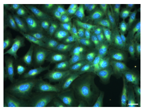 | BRD-K47520858-001-02-6 Reference Molecule | MGMG 1.00 | BioT5 0.58 | MolT5 0.22 | CPMolGAN 0.18 |
| The molecule is a urea in which one of the nitrogens is substituted by a 4-(trifluoromethyl)phenyl group, while the other is substituted by a tetrahydropyran-3-yl group. It is a synthetic oral anticoagulant that targets activated factor Xa in the coagulation cascade. It has a role as an anticoagulant, an EC 3.4.21.6 (coagulation factor Xa) inhibitor and a serine protease inhibitor. It contains a morpholine, a secondary amide, and a primary alcohol. | 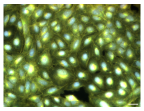 | BRD-K61494613-001-01-3 Reference Molecule | MGMG 1.00 | BioT5 0.45 | MolT5 0.16 | CPMolGAN 0.12 |
| The molecule is a member of the class of ureas that is urea in which the hydrogens of the primary amino group have been replaced by methyl groups, while the hydrogens attached to the nitrogen are replaced by a 1-hydroxypropan-2-yl group and a hydrogen attached to the other nitrogen is replaced by a (2S,3R)-8-(2-fluorophenyl)-5-[(2R)-1-hydroxypropan-2-yl]-3-methyl-6-oxo-2H,3H,4H,5H,6H-pyrido[2,3-b][1,5]oxazocin-2-yl]methyl}. It is an ether, a member of ureas, a member of monofluorobenzenes, and a monofluorophenyl. | 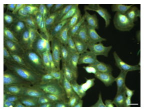 | BRD-K97106314-001-01-3 Reference Molecule | MGMG 0.82 | BioT5 0.26 | MolT5 0.28 | CPMolGAN 0.17 |
| The molecule is a pyridine carboxamide obtained by formal condensation of the carboxy group of 4-nicotinic acid with the amino group of (2S,3R)-2-[(cyclopropylmethyl)(methyl)amino]-1-hydroxypropyl 3-(methylamino)-2,3-dihydro-1H-indol-5-one. This contains a pyridine carboxamide, a tertiary amino compound, an aromatic amide and a monocarboxylic acid amide. | 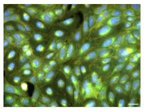 | BRD-K93819193-001-01-1 Reference Molecule | MGMG 0.85 | BioT5 0.65 | MolT5 0.34 | CPMolGAN 0.14 |

**Supplementary Fig. 2: Example comparisons of the top 1 molecule generated by MGMG and unimodal baselines based on the same textual descriptions and morphological profiles of reference small molecules.**

| Textual Description | Morphological Profile | Output | | |
|---|---|---|---|---|
| The molecule is a member of the class of phenylureas that is urea in which one of the nitrogens is substituted by a phenyl group, while the other is substituted by a 4-chlorophenyl group. It contains a phenylurea, monochlorobenzene, an ether and a tertiary amine. | 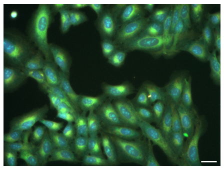 | BRD-K99353774-001-01-2 Reference Molecule | Sample #1 0.86 | Sample #2 0.81 |
| | | Sample #3 0.81 | Sample #4 0.69 | Sample #5 0.69 |
| The molecule is a azetidine-2-carbonitrile that is substituted at positions 1, 2, 3 and 4 by 1-(pyridine-3-carbonyl) azetidine, 2-carbonitrile, 3-{4-[2-(2-fluorophenyl)ethynyl]phenyl}, and 4-(hydroxymethyl), respectively (the 2S,3R,4S, diastereoisomer). This contains an azetidine, a primary alcohol, monofluorinated phenyl, nitrile, and pyridine. | 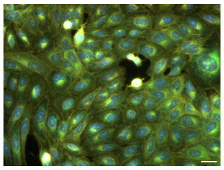 | BRD-K07799833-004-02-8 Reference Molecule | Sample #1 1.00 | Sample #2 1.00 |
| | | Sample #3 0.81 | Sample #4 0.61 | Sample #5 0.61 |
| The molecule is a member of the class of azabicycloalkanes. It has a role as a protein kinase inhibitor. It contains a monofluorobenzene, a 2-pyridone, a piperazine, a cyclohexane, a primary alcohol,and a monocarboxylic acid. | 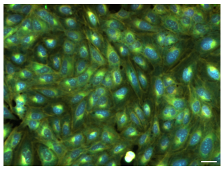 | BRD-K15460493-001-01-6 Reference Molecule | Sample #1 0.81 | Sample #2 0.81 |
| | | Sample #3 0.70 | Sample #4 0.63 | Sample #5 0.54 |
| The molecule is alpha-Ergocryptine with a bromine atom added to the ergoline scaffold. It is a natural ergot alkaloid. Note that ergocryptine discussed in the literature prior to 1967, when beta-ergocryptine was separated from alpha-ergocryptine, is now referred to as alpha-ergocryptine. | 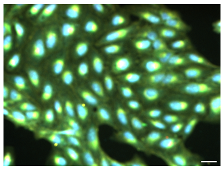 | BRD-A60274948-066-02-9 Reference Molecule | Sample #1 1.00 | Sample #2 0.86 |
| | | Sample #3 0.83 | Sample #4 0.41 | Sample #5 0.40 |

**Supplementary Fig. 3: Examples of the top 5 molecules generated by MGMG based on the textual description and morphological profile of a small molecule.**

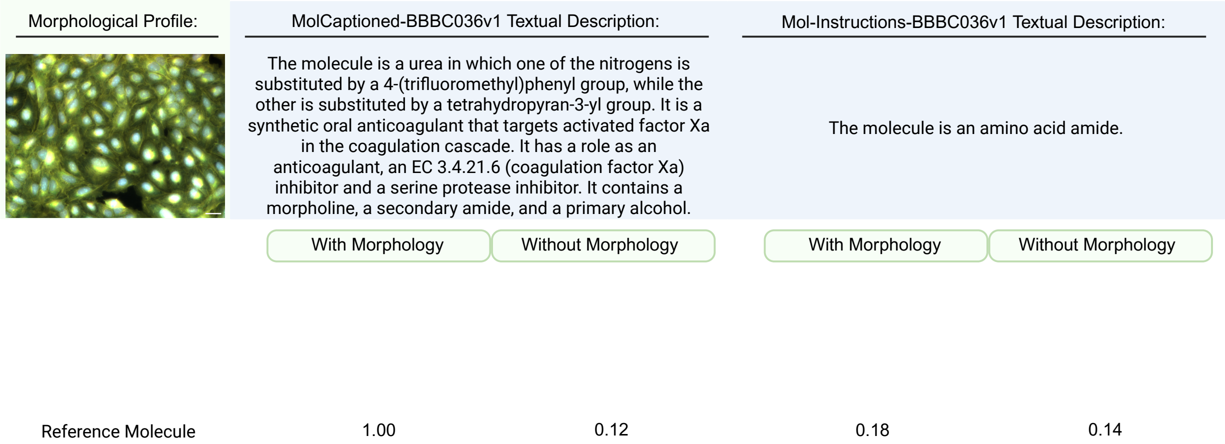


**Supplementary Fig. 4. An example of molecules generated by MGMG using the same morphological profile but paired with either rich-informative or under-informative textual descriptions from the MolCaptioned-BBBC036v1 and Mol-Instructions-BBBC036v1 datasets.**

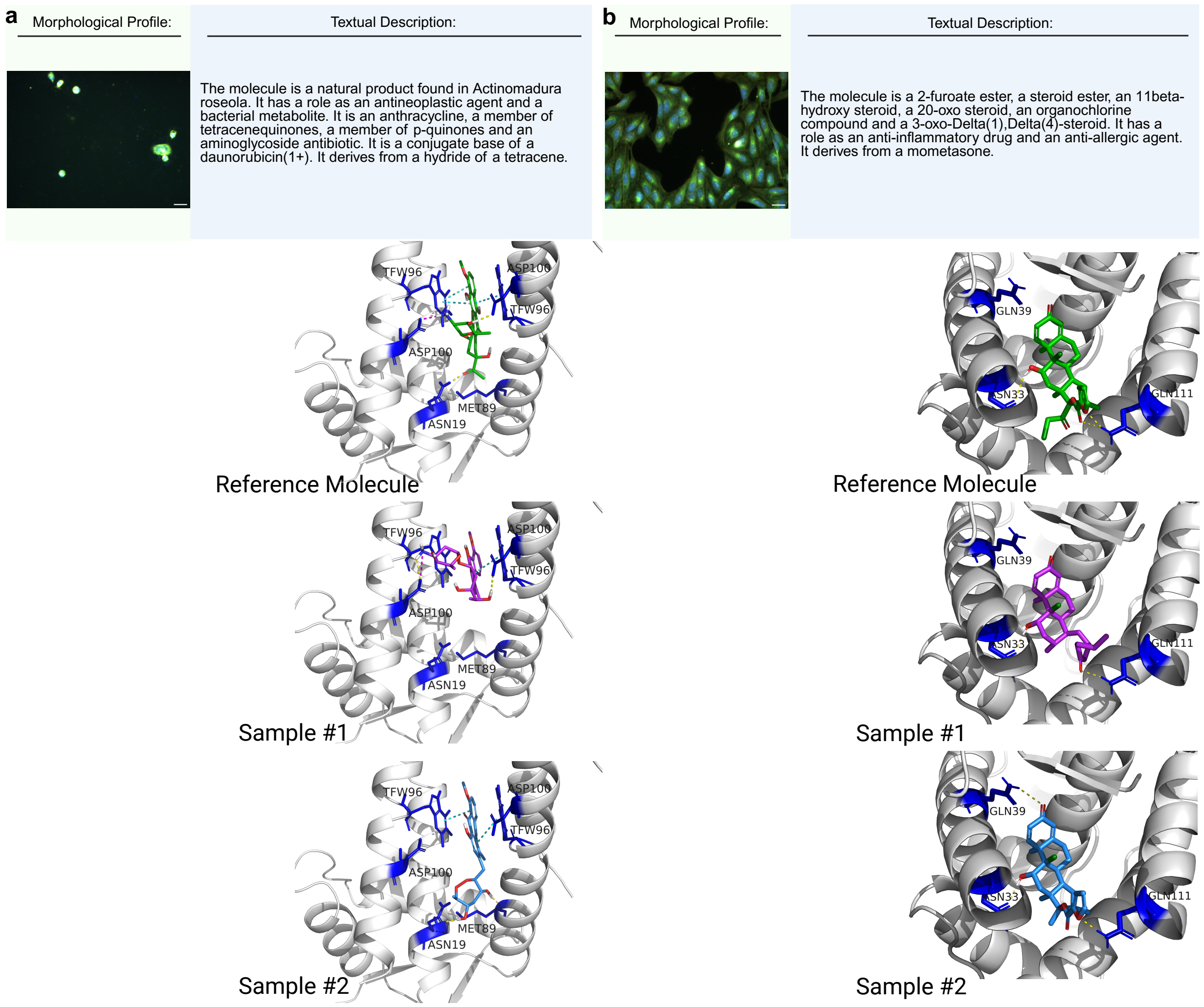


| PDB ID: 7QZ7 | Reference | Sample #1 | Sample #2 |
|---|---|---|---|
| Similarity to reference | - | 0.746 | 0.424 |
| Docking score | -10.45 | -8.54 | -10.72 |
| QED | 0.305 | 0.335 | 0.543 |
| SA | 5.410 | 4.676 | 4.274 |

| PDB ID: 4E2J | Reference | Sample #1 | Sample #2 |
|---|---|---|---|
| Similarity to reference | - | 0.384 | 0.786 |
| Docking score | -11.54 | -10.19 | -12.00 |
| QED | 0.449 | 0.610 | 0.494 |
| SA | 4.834 | 5.212 | 4.845 |

**Supplementary Fig. 5: Additional examples of molecular docking results of MGMG generated molecules.**
